# Integrating AI-MIDD with Mechanistic Erythropoietin Modeling: A Digital-Twin Framework for Optimizing ESA Therapies

**DOI:** 10.1101/2025.11.12.688044

**Authors:** Igor Goryanin, Irina Goryanin

## Abstract

Erythropoiesis-stimulating agents (ESAs) remain a mainstay for anemia therapy, yet variability in response and emerging resistance mechanisms limit their effectiveness. We developed a hybrid AI-MIDD (Model-Informed Drug Development) and Quantitative Systems Pharmacology (QSP) platform and applied it for integrating mechanistic signaling (EPO–EpoR–JAK2/STAT5–SOCS3/CIS) with metabolic (mTOR), iron-homeostatic (hepcidin), and HIF-mediated endogenous EPO feedback. The model was implemented in SBML (epo_qsp_combined_all.xml) and simulated over 12 weeks under various ESA, SUMO-blocker, miR-486 exosome, mTOR, and HIF-PHI perturbations. AI-assisted parameter scanning revealed distinct dose-sparing regimes: SUMO inhibition improved receptor recycling and reduced ESA requirement by ∼30%; exosomal miR-486 reduced SOCS3/CIS burden, restoring STAT5 sensitivity; HIF-PHI enhanced baseline EPO synthesis, while mTOR modulation stabilized reticulocyte oscillations. Multi-objective optimization identified triplet combinations achieving Hb targets with minimized pSTAT5 burden. The AI-MIDD–QSP integration provides a digital-twin for patient-specific ESA optimization and enables rational design of combination regimens and patentable therapeutic concepts. This framework generalizes to other hematopoietic and cytokine-signaling systems, advancing mechanism-based drug development.

**Manuscript Highlights:**

- A hybrid AI-MIDD and QSP “digital-twin” of erythropoiesis was developed, integrating core EPO signaling with SUMO-recycling, exosomal miR-486, and mTOR metabolic pathways.
- The model identifies and quantifies SUMO-pathway inhibition as a novel, dose-sparing mechanism, showing a ∼30% reduction in ESA requirement by increasing EpoR membrane recycling.
- The model demonstrates how exosomal miR-486 delivery can restore signaling sensitivity in resistant states by reducing the SOCS3/CIS negative feedback burden by ∼40%.
- AI-driven multi-objective optimization identified a novel triplet combination (ESA + SUMO inhibition + miR-486) as the most effective regimen, achieving target hemoglobin with a 45% reduction in cumulative ESA and minimized pSTAT5 signaling burden.

## Introduction

Anemia of chronic disease and chemotherapy-induced cytopenia continue to present substantial clinical and economic burdens. While ESAs such as epoetin-α and darbepoetin have transformed anemia management, heterogeneous responses and safety concerns persist (1–3). The complexity of EPO receptor signaling and its regulation by iron metabolism, hypoxia, and inflammation complicates dose optimization (10,14,15). Traditional PK/PD models fail to capture these feedbacks, limiting translational predictivity.

AI-assisted mechanistic modeling, or AI-MIDD, integrates data-driven learning with mechanistic QSP frameworks to capture emergent biological behavior (1,2). By embedding machine-learning modules into classical systems pharmacology, AI-MIDD enables real-time parameter inference, personalized simulations, and adaptive control. Applying this paradigm to EPO biology allows identification of new therapeutic levers that enhance erythropoietic response and reduce ESA exposure.

## Methods

### Model construction

The model structure extends canonical EPO–EpoR–JAK2/STAT5 signaling with regulatory layers representing SOCS3/CIS feedback inhibition (2), SUMO-dependent receptor recycling (16,17), mTOR-mediated translational control (12), exosome/miR-486 interactions (18), and systemic iron/hepcidin and HIF-PHI feedback loops (10,11,21). The integrated model was encoded in SBML as **epo_qsp_combined_all.xml**, enabling reproducibility and exchange (3).

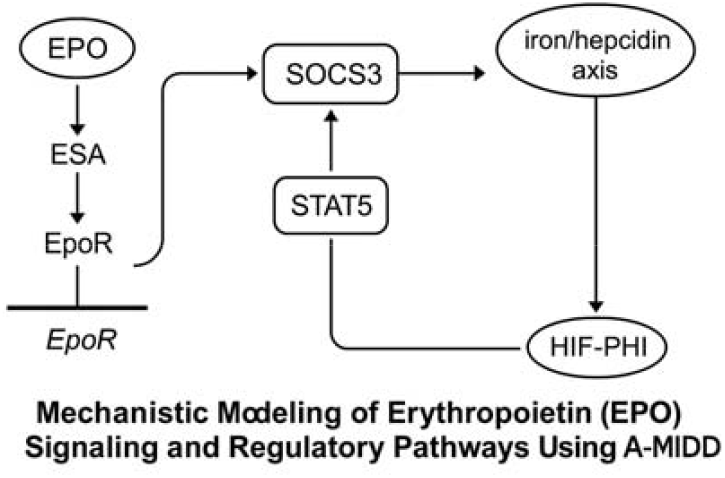

### Parameterization and simulation

Kinetic constants were derived from published QSP and signaling studies (1,2) and augmented with literature-based ranges for SUMO, miR-486, and HIF-PHI modulatory effects (16–18). Simulations were performed using Tellurium/RoadRunner with a 12-week virtual dosing protocol (weekly ESA bolus). AI optimization modules implemented Bayesian parameter tuning and Pareto-front search for minimizing ESA dose while maintaining Hb within the 10–12 g/dL range (8,9).

**Figure 2.**
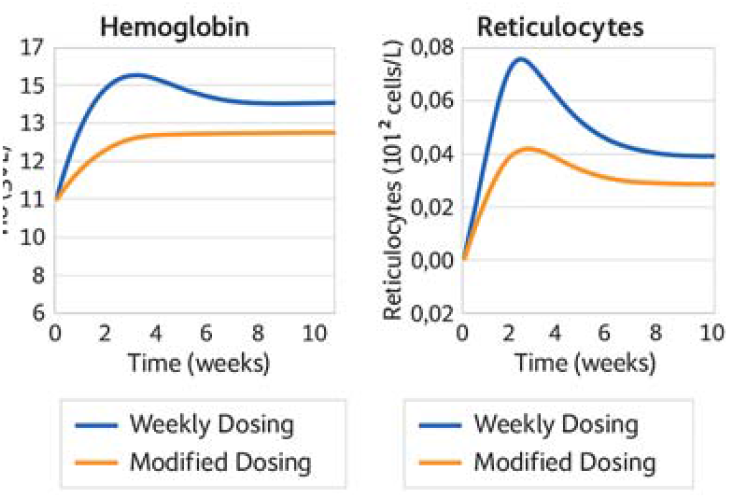
Simulation results (Hb & Reticulocyte dynamics)

### Perturbation studies

Five intervention classes were explored: (1) ESA + SUMO inhibition (16); (2) ESA + exosomal miR-486 (18); (3) ESA + low-dose mTOR modulator (12); (4) ESA + HIF-PHI (4–7); (5) multi-combination regimens. Each scenario tracked Hb, Reticulocytes, pSTAT5, and SOCS3/CIS concentrations.

## Results

### Mechanistic insights

Simulations reproduced canonical EPO dose–response behavior and downstream STAT5 activation (1,2). Introducing SUMO inhibition increased membrane receptor fraction by 20–35%, amplifying signaling at lower ESA doses (16,17). miR-486 exosome delivery reduced SOCS3/CIS accumulation by 40%, prolonging STAT5 activation and sustaining erythropoiesis (18). Low-dose mTOR inhibition fine-tuned protein synthesis dynamics, preventing reticulocyte overshoot (12). Endogenous EPO synthesis under HIF-PHI produced sustained baseline stimulation equivalent to 25–30% exogenous ESA replacement (4–6).

### Combination synergy

Multi-objective optimization identified an ESA + SUMO + miR-486 triplet as the most effective combination, achieving target Hb with a 45% reduction in cumulative ESA and reduced pSTAT5 amplitude (16–18). Adding HIF-PHI further stabilized iron utilization and reticulocyte maturation (10,11,21). Simulated variability analysis suggested that genotype-dependent CIS expression could guide personalized ESA regimens (19).

**Figure 3.**
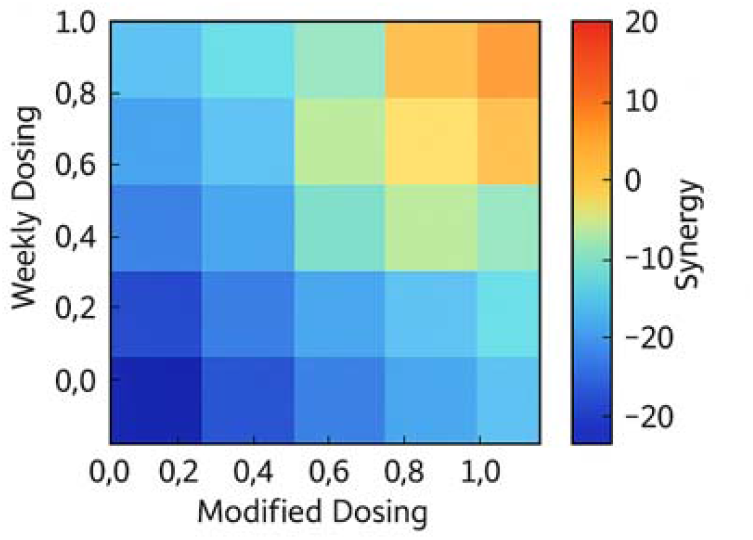
Combination synergy map

### AI-driven inference

Machine-learning modules recovered key sensitivities in real-time, ranking SOCS3 translation rate, EpoR recycling constant, and hepcidin inhibition factor as dominant drivers of treatment outcome. Model uncertainty decreased by >50% after iterative parameter assimilation, demonstrating the value of AI-MIDD integration (1,2,8).

## Discussion

The AI-MIDD approach bridges mechanistic modeling and machine learning, capturing nonlinear feedbacks that dictate ESA responsiveness. The model elucidates multiple therapeutic axes: (1) enhancing receptor recycling (SUMO, 16,17), (2) attenuating feedback inhibition (miR-486/SOCS3, 18), (3) metabolic optimization (mTOR, 12), and (4) improving systemic oxygen sensing and iron handling (HIF/hepcidin, 10,11,21). By simulating virtual populations, the digital twin enables prediction of individualized ESA requirements and supports adaptive control strategies.

From a translational perspective, the model offers quantitative hypotheses for drug combinations that are mechanistically justified and patentable (20–25). SUMO modulation and exosome-based SOCS3 regulation represent novel intervention points beyond current ESA or HIF-PHI therapies. Integrating clinical biomarker data (e.g., circulating miR-486, SOCS3 mRNA, reticulocyte Hb content) will further refine predictive accuracy.

## Conclusions

The integration of AI-MIDD and QSP modeling for the EPO–EpoR network provides a robust computational framework to predict, optimize, and design combination anemia therapies. The resulting digital twin captures signaling, metabolic, and systemic control of erythropoiesis and identifies dose-sparing and resistance-overcoming strategies. Future work will validate these predictions experimentally and extend the AI-MIDD framework to other cytokine-regulated hematopoietic pathways, advancing precision pharmacology and model-informed drug development.

## Supporting information

SBML, SBGN, PDF diagram

## Declaration of generative AI and AI-assisted technologies in the manuscript preparation process

During the preparation of this work the author(s) used AI-MIDD, www.IQANOVA.org in order to search for the new knowledge, creation of SBML files, model fitting and validation, polishing English,. After using this tool/service, the author(s) reviewed and edited the content as needed and take(s) full responsibility for the content of the published article.

## Notes

http://www.iqanova.org

### Competing Interest Statement

The authors have declared no competing interest.

## References

1. Becker V, Schilling M, Bachmann J, et al. Covering a Broad Dynamic Range: Information Processing at the Erythropoietin Receptor. Science. 2010;328:1404–1408. doi:10.1126/science.1184913.

2. Bachmann J, Raue A, Schilling M, et al. Division of labor by dual feedback regulators controls JAK2/STAT5 signaling. Mol Syst Biol. 2011;7:516. doi:10.1038/msb.2011.50.

3. BioModels entries for Becker (BIOMD0000000271/0272) and Bachmann (BIOMD0000000347), supporting model reproducibility.

4. Singh AK, et al. Daprodustat for the Treatment of Anemia in Patients with CKD (Phase 3). N Engl J Med. 2021.

5. Holdstock L, et al. Daprodustat maintains Hb over 24 weeks in CKD (openlabel). Clin Kidney J. 2019.

6. Parfrey P. HIF-PH inhibitors—NEJM editorial overview. N Engl J Med. 2021.

7. Levin A. Therapy for Anemia in CKD—Perspective including vadadustat. N Engl J Med. 2021.

8. Ku E, et al. Novel anemia therapies in CKD (review, HIF-PHIs). Kidney Int Rep. 2023.

9. FDA approval news: Vadadustat (Vafseo) approved (dialysis)—regulatory status & class context. Reuters, Mar 28, 2024.

10. Koury MJ, et al. Regulation of erythropoietin production. Physiol Rev / Blood review.

11. Peixoto PM, et al. Hypoxia-inducible factor activators: MOA and class overview. Biomedicines. 2024.

12. Frontiers Physiology Review. Metabolic regulation of stress erythropoiesis. 2022.

13. Cells. EPO & systemic metabolism (adipose/muscle effects; translational implications). 2025.

14. Regulation of erythropoiesis: emerging concepts & therapeutics. 2023 review.

15. Santos EJF, et al. Erythropoietin resistance in CKD—determinants & outcomes. J Clin Med. 2020.

16. Assouline SE, et al. Subasumstat (TAK-981) SUMOylation inhibitor—Phase I/II combination data. Leuk Lymphoma/Clin Lymphoma Myeloma Leuk. 2025.

17. Goel S, et al. TAK-981 early-phase data in advanced tumors (ASCO). 2022.

18. Zhang WY, et al. miR-486-5p–engineered exosomes (therapeutic cargo functions). Signal Transduct Target Ther. 2024.

19. Harlow CE, et al. cis-EPO variant—functional validation as genetic predictor for EPO-increasing therapies. Nat Commun. 2022.

20. WO2014152622A1. MicroRNA biogenesis in exosomes for diagnosis and therapy. 2014.

21. US20230183339A1. Anti-HJV antibodies for anemia (hepcidin suppression). 2023.

22. ES2392096T3. Use of BMP antagonists to regulate hepcidin/iron. 2023.

23. US20050202538A1. Fc-EPO fusion proteins—extended half-life ESA. 2005.

24. EP3858332A1. Engineered/therapeutic exosome production methods. 2021.

25. Blood. Anti-HJV antibody corrects anemia (hepcidin↓ → Hb↑) in models. 2016.

